# Equivalence of LD-Score Regression and Individual-Level-Data Methods

**DOI:** 10.1101/211821

**Authors:** Ronald de Vlaming, Magnus Johannesson, Patrik K.E. Magnusson, M. Arfan Ikram, Peter M. Visscher

## Abstract

LD-score (LDSC) regression disentangles the contribution of polygenic signal, in terms of SNP-based heritability, and population stratification, in terms of a so-called intercept, to GWAS test statistics. Whereas LDSC regression uses summary statistics, methods like Haseman-Elston (HE) regression and genomic-relatedness-matrix (GRM) restricted maximum likelihood infer parameters such as SNP-based heritability from individual-level data directly. Therefore, these two types of methods are typically considered to be profoundly different. Nevertheless, recent work has revealed that LDSC and HE regression yield near-identical SNP-based heritability estimates when confounding stratification is absent. We now extend the equivalence; under the stratification assumed by LDSC regression, we show that the intercept can be estimated from individual-level data by transforming the coefficients of a regression of the phenotype on the leading principal components from the GRM. Using simulations, considering various degrees and forms of population stratification, we find that intercept estimates obtained from individual-level data are nearly equivalent to estimates from LDSC regression (*R*^2^ *>* 99%). An empirical application corroborates these findings. Hence, LDSC regression is not profoundly different from methods using individual-level data; parameters that are identified by LDSC regression are also identified by methods using individual-level data. In addition, our results indicate that, under strong stratification, there is misattribution of stratification to the slope of LDSC regression, inflating estimates of SNP-based heritability from LDSC regression *ceteris paribus*. Hence, the intercept is not a panacea for population stratification. Consequently, LDSC-regression estimates should be interpreted with caution, especially when the intercept estimate is significantly greater than one.

---

Population stratification can confound genome-wide association study (GWAS) summary statistics, as stratification may inflate *χ*^2^-test statistics^1–4^. LD-score (LDSC) regression incorporates a parameter, referred to as the ‘intercept’, that accounts at least partially for confounding stratification in GWAS summary statistics^4^. By also including linkage-disequilibrium (LD) scores as a regressor, this method is able to disentangle the contribution of stratification and polygenic signal to GWAS test statistics.

Stratification can also bias estimates of variance components^5^ in a linear mixed model (LMM), as admixture affects the inferred genetic relatedness between individuals^6,7^ and, thereby, the relatedness matrix and its eigenvalues^8^. By including the leading principal components (PCs) from the genomic-relatedness matrix (GRM; inferred e.g., using GCTA^9^ or PLINK^10,11^) as fixed-effect covariates in genomic-relatedness-matrix restricted maximum likelihood (GREML) estimation, one can correct for the confounding effects of stratification on the inferred variance components^5^.

At first glance, LDSC regression and individual-level-data methods seem weakly related; both LDSC regression and GREML estimation can be used to infer SNP-based heritability 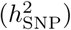 and both methods account for LD^4,12^. In fact, in case population stratification is absent, LDSC and Haseman-Elston (HE) regression^13^ are essentially equivalent when estimating 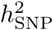 ^14^. Importantly, HE regression is a simplified form of GREML estimation. The relation between LDSC regression and individual-level data methods, however, seems to break down when considering the intercept in LDSC regression. An equivalent parameter is not reported by methods such as GREML estimation. Nevertheless, as both approaches assume the same data-generating process, we assert that the equivalence can be extended to include the intercept.

## Estimating the LD-Score-Regression Intercept from Individual-Level Data Directly

We ascertain this thesis by studying how the GRM behaves when generalizing the framework for population stratification assumed in LDSC regression^4^ to multiple discrete subpopulations. Within this framework, we derive an ‘expected’ GRM and an explicit eigendecomposition of this matrix for *P* = 2 discrete subpopulations, in line with LDSC regression. In this case, the leading PC and phenotype vector can be used to infer the LDSC intercept directly. Specifically, like the LDSC framework, we (i) consider a pooled sample with no close relatives, with *n* individuals from both subpopulations yielding a pooled sample size *N* = 2*n*, (ii) assume the phenotype is standardized, and (iii) assume differences in allele frequencies are shaped by Wright's *F*-statistic^4,15,16^ (*F*_ST_). We show that this genetic drift induces subtle negative ‘relatedness’ between populations and positive ‘relatedness’ within populations. An individual-level-data estimator of the intercept in this scenario is given by

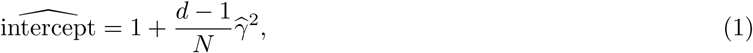

where 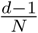 is an estimate of *F*_ST_, based on the leading EV (*d*) of the GRM, and where 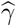 is the estimate of a linear regression of the standardized phenotype on the leading PC. Formal derivations are reported in Appendix A. In general, the ordinary least squares (OLS) and generalized least squares (GLS) estimator of *γ* are not identical. However, in this particular model these estimators are equivalent. In addition to OLS and GLS regression, we can estimate the intercept by extending an HE regression^13^. An overview of the methods is shown in Table 1. For more complex forms of stratification with *P* subpopulations, we posit that

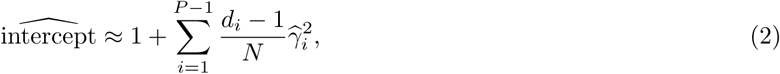

**Table 1.**
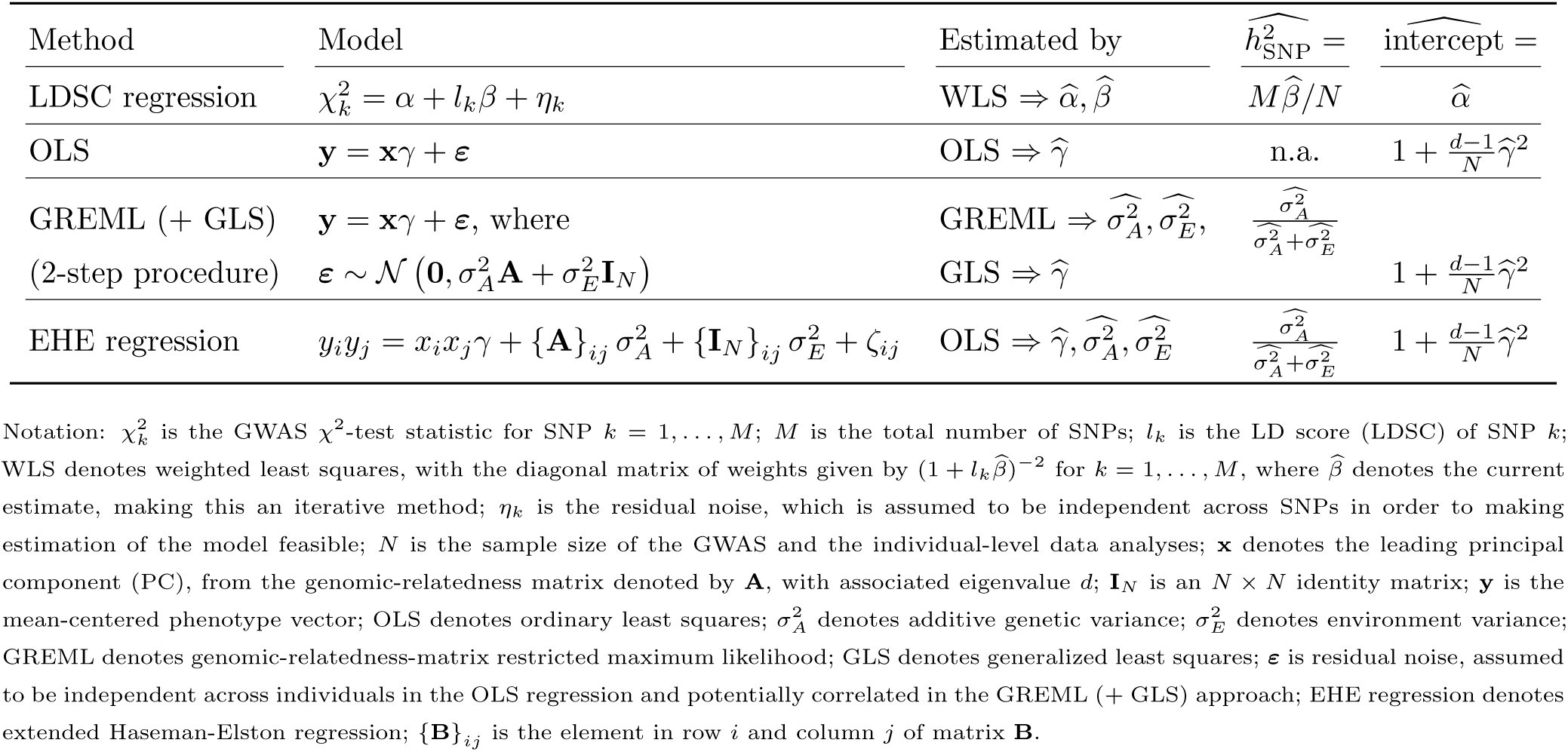
Methods for jointly estimating the SNP-based heritability 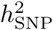 and the LD-score intercept, the latter parameter reflecting the amount of confounding stratification present for a phenotype in a given sample.

where *N* is the pooled sample size and where *d*_*i*_ denotes the *i*-th leading eigenvalue of the GRM and 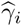 the estimate of a linear regression of the standardized phenotype on the *i*-th PC.

## Simulations and Empirical Analyses Using Data Exhibiting Population Structure

We assess the accuracy of Equations 1 and 2 by means of two sets of simulations, based on pooled genotype data from the Swedish Twin Registry (STR), the Health and Retirement Study (HRS), and the Rotterdam Study (RS)^17^. Details on the quality control (QC) of these data are reported in Appendix B. After QC we have *N* = 17, 544 observations (*n* = 5, 848 from each of the three subsamples) and *M* = 1, 023, 716 HapMap 3 SNP_s_^18^ with minor allele frequency greater than 1%. In addition, we perform empirical analyses, to assess the merits of Equation 2 in real data, where subtle stratification may be at play and where the assumed discrete nature of stratification, with equal sample size per subpopulation, may break down.

Figure 1 shows the scatter plot of the leading two principal components of the GRM. There is clear clustering of the STR, HRS, and RS samples. Although the individuals from these studies are not fully separated along the first and second PC, the separation is quite accurate; when classifying the lower-left quadrant as HRS, the upper-left quadrant as RS, and the right half as STR, 92% of the individuals are correctly classified. Regardless of the etiology of this clustering (e.g., differences in true allele frequencies and batch effects), the clustering shows that we have a dataset that closely follows the theoretical assumptions of LDSC regression.

**Figure 1.**
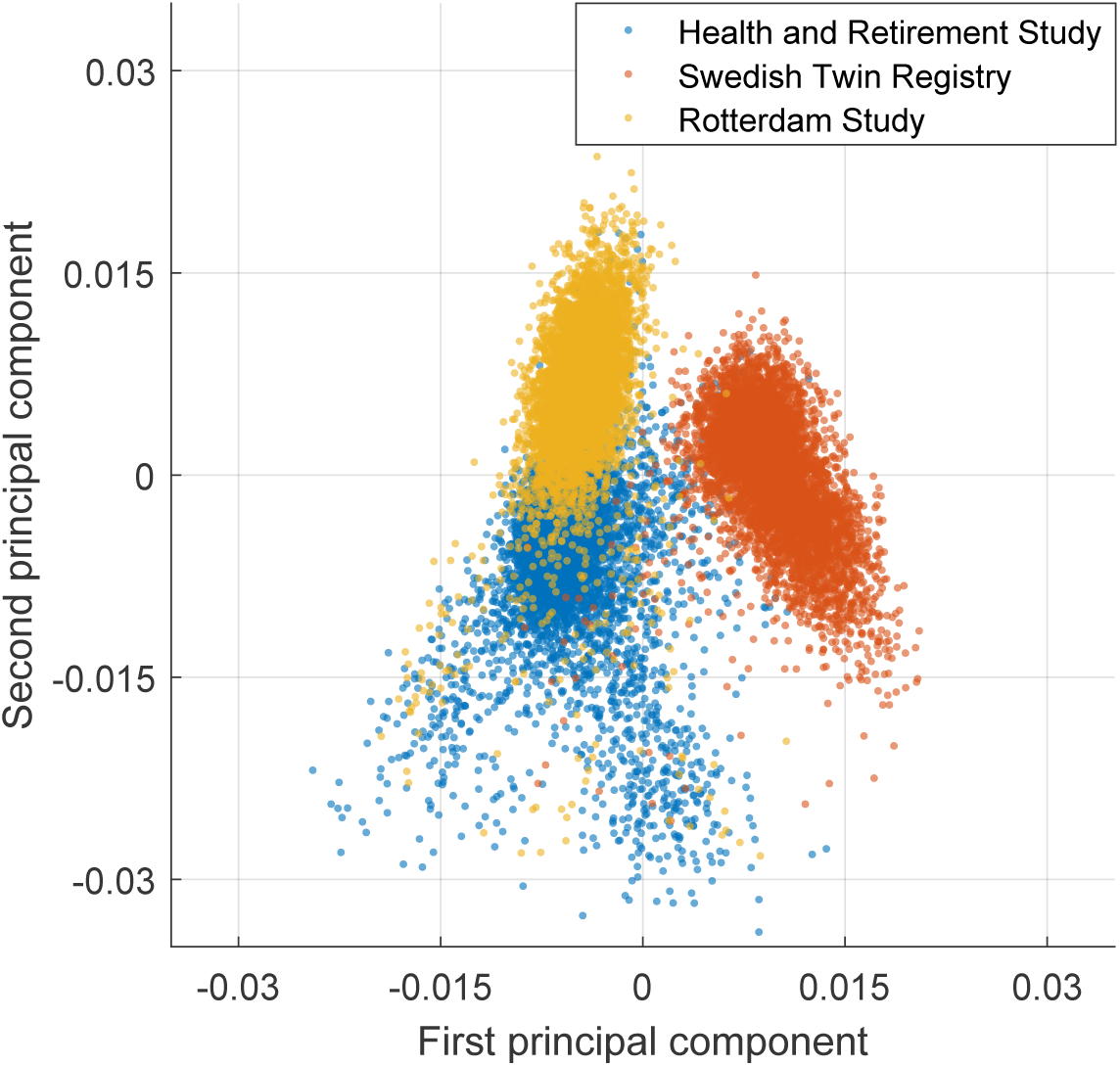
Scatter plot of the leading two principal components from the genomic-relatedness matrix of pooled data from the Health and Retirement Study (in blue), the Swedish Twin Registry (orange), and the Rotterdam Study (yellow).

In all simulations, we use this genetic data to simulate phenotypes having a (i) polygenic architecture and (ii) difference in phenotypic mean between the different subsamples. We apply GREML (followed by GLS or OLS) to the simulated data, to estimate the intercept and 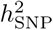 from individual-level data, and use LDSC regression to estimate the same parameters using GWAS results from the same samples. We compare resulting estimates. In addition, in each simulation we compute the attenuation ratio^19^, defined as the LDSC-regression intercept estimate minus one and the average *χ*^2^-test statistic across markers 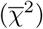 minus one. For this ratio it holds that

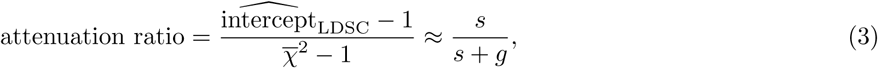

where *s* reflects inflation due to stratification and *g* inflation due to signal. Hence, this ratio quantifies the proportion of inflation in GWAS *χ*^2^-test statistics, away from one, that can be attributed to stratification.

We consider five levels of stratification, explaining from 0% up to 20% of the phenotypic variance. The stratification is shaped by differences in phenotypic mean between the HRS, STR, and RS samples. For each level of stratification we simulate 500 phenotypes. Regarding polygenic architecture, each phenotype follows an infinitesimal model, where each SNP is standardized and where standardized SNP_s_ have effects that are normally distributed and independent draws, with SNP-effect sizes and residual variance such that 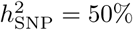. The simulation design is discussed in detail in Appendix C. As GREML estimation is computationally expensive, we derive an efficient GREML algorithm in Appendix D. This algorithm can be used when only PCs are included as fixed-effect covariates.

In the first set of simulations, we gauge the accuracy of Equation 1 (i.e., for *P* = 2 subpopulations) by considering only the STR and HRS samples. In the second simulation, we assess the accuracy of our extrapolation in Equation 2, by considering the full sample (i.e., *P* = 3). In the empirical analyses, we also assess the accuracy of Equation 2 using human height and body-mass index (BMI) as outcomes. As these phenotypes are first standardized at the study level, we assume there is only subtle stratification at play. Hence, we set *P* relatively high (i.e., *P* = 20). In a semi-empirical extension, we introduce artificial stratification by assigning the HRS, STR, and RS samples different phenotypic means, keeping *P* = 20. In all analyses, we apply LDSC regression and GREML estimation.

Finally, we consider an additional set of simulations for the HRS and STR samples, where we compare different sources of stratification and different means to control for it. More specifically, we simulate data where stratification is either shaped by the lead PC as inferred from the GRM or by differences in mean between subsamples, where in both cases stratification explains 20% of the phenotypic variance. For both scenarios, we assess how LDSC behaves when (i) failing to control for stratification in the GWAS, (ii) when controlling for it using a subsample dummy, and (iii) using the lead PC. Similarly, we assess the behavior of GREML when using either the subsample dummy or the lead PC as fixed-effect covariate for estimating the intercept and controlling for stratification when estimating 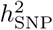

## Results for Two Discrete Populations

Figure 2 shows intercept and 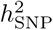 estimates from LDSC regression and the GREML for 500 independent runs and for various levels of stratification. Across the runs and levels of stratification, the intercept estimates are of the same scale and highly correlated (*R*^2^ = 99.89%). For 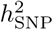 the results diverge; the estimates across runs and levels of stratification are weakly correlated (*R*^2^ = 18.17%). However, more importantly, as can be seen in Panel A of Table 2, there is a strong increase in 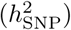 estimates from LDSC regression as the amount of stratification increases. For the design with no stratification, the average 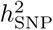 estimate of LDSC regression is ~ 51% whereas in the design with the highest amount of stratification, the average estimate is ~ 94%. Hence, in relative terms, the LDSC-regression 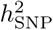 estimate under strong stratification is ~ 84% higher than the estimate under no stratification, while in both instances in truth 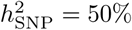. Although with GREML estimation, controlling for the first PC (inferred empirically from the GRM), we also see an increase with the amount of stratification, this increase is far smaller; under no stratification the average estimate is ~ 50%, while for the highest amount of stratification the estimate is ~ 54%.

**Figure 2.**
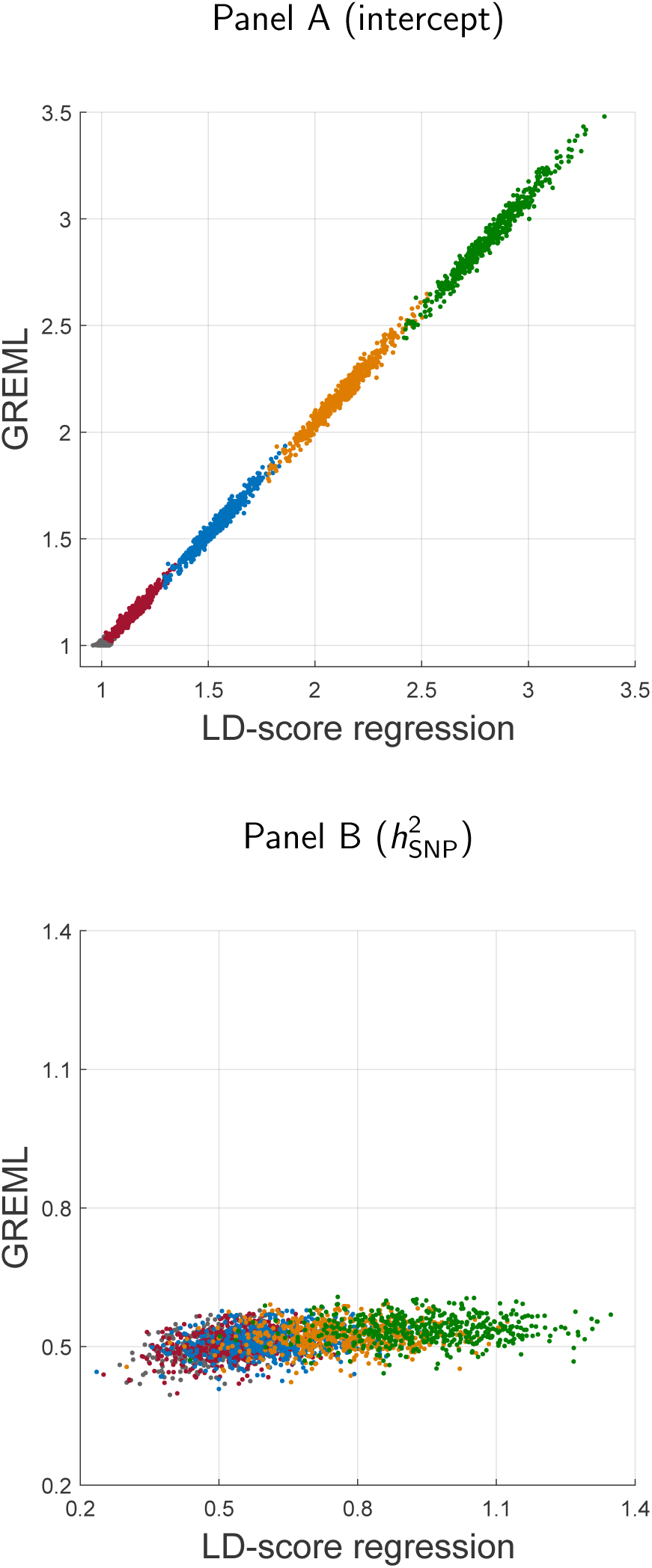
Scatter plots of estimates of the LD-score-regression intercept (Panel A) and SNP-based heritability (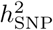; Panel B) across 500 runs and various levels of stratification based on data from *P* = 2 subpopulations. *x*-axis: LD-score regression estimates. *y*-axis: GREML estimates. In the GREML approach, the leading principal component is used as fixed-effect covariate, and the estimate of that fixed effect 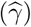 is cast to an intercept estimate using Equation 1. Gray dots: no stratification; red dots: light stratification; blue dots: moderate stratification; yellow dots: substantial stratification; green dots: strong stratification.

**Table 2.** Mean of estimates of SNP-based heritability (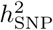), intercept, and the attenuation ratio, across 500 runs and corresponding standard errors (s.e.), for GREML estimation and LDSC regression, for various levels of stratification, for pooled data from *P* = 2 subpopulations (i.e., STR and HRS; Panel A) and *P* = 3 subpopulations (i.e., STR, HRS, and RS; Panel B).

| Panel A: $P = 2$ | | | | | |
| --- | --- | --- | --- | --- | --- |
| Method | None | Light | Stratification:<br>Moderate | Substantial | Strong |
| Mean $h_{\text{SNP}}^2$ ( s.e. ) | | | | | |
| LDSC | 50.65% ( 0.35% ) | 53.58% ( 0.39% ) | 62.57% ( 0.48% ) | 76.65% ( 0.58% ) | 94.28% ( 0.66% ) |
| GREML | 50.28% ( 0.13% ) | 50.51% ( 0.13% ) | 51.19% ( 0.13% ) | 52.28% ( 0.13% ) | 53.72% ( 0.13% ) |
| Mean intercept ( s.e. ) |  |  |  |  |  |
| LDSC | 1.011 ( 0.001 ) | 1.149 ( 0.003 ) | 1.543 ( 0.005 ) | 2.128 ( 0.006 ) | 2.825 ( 0.007 ) |
| GREML | 1.007 ( 0.000 ) | 1.152 ( 0.003 ) | 1.566 ( 0.005 ) | 2.183 ( 0.007 ) | 2.918 ( 0.008 ) |
| Mean attenuation ratio ( s.e. ) |  |  |  |  |  |
| LDSC | 0.086 ( 0.006 ) | 0.531 ( 0.005 ) | 0.790 ( 0.002 ) | 0.867 ( 0.001 ) | 0.896 ( 0.001 ) |
| Panel B: $P = 3$ | | | | | |
| Method | None | Light | Stratification:<br>Moderate | Substantial | Strong |
| Mean $h_{\text{SNP}}^2$ ( s.e. ) | | | | | |
| LDSC | 50.59% ( 0.24% ) | 51.69% ( 0.27% ) | 54.63% ( 0.39% ) | 59.02% ( 0.60% ) | 64.35% ( 0.87% ) |
| GREML | 49.99% ( 0.08% ) | 50.52% ( 0.08% ) | 52.01% ( 0.09% ) | 54.26% ( 0.11% ) | 57.00% ( 0.15% ) |
| Mean intercept ( s.e. ) |  |  |  |  |  |
| LDSC | 1.012 ( 0.001 ) | 1.122 ( 0.003 ) | 1.427 ( 0.008 ) | 1.878 ( 0.016 ) | 2.413 ( 0.025 ) |
| GREML | 1.008 ( 0.000 ) | 1.117 ( 0.003 ) | 1.417 ( 0.009 ) | 1.862 ( 0.019 ) | 2.390 ( 0.030 ) |
| Mean attenuation ratio ( s.e. ) |  |  |  |  |  |
| LDSC | 0.064 ( 0.004 ) | 0.388 ( 0.006 ) | 0.670 ( 0.004 ) | 0.797 ( 0.003 ) | 0.858 ( 0.001 ) |

As the LDSC-regression framework assumes allele-frequency differences, when standardized by pooled frequency, are homoskedastic random draws with mean zero and a variance equal to *F*_ST_, we investigate whether the allele-frequency differences satisfy this assumption. Figure 3 shows a histogram of these differences, when setting the coding allele randomly. The mean of differences is − 1.13 ×10^*−*5^ and the variance is 9.2 ×10^*−*4^, which is close to *F*_ST_ as estimated from the leading eigenvalue (*d*_1_), *viz*., *F*_ST_ = *N ^−^*^1^ (*d*_1_), *viz.*, *F*_ST_ =(*d*_1_ − 1) = 9.8 ×10^*−*4^. Moreover, these differences seem normally distributed. Nevertheless, the Jarque-Bera test for normality^20^ rejects the null, with a test statistic of 318. Hence, these differences are statistically non-normally distributed. However, when excluding the 49 SNP_s_ with allele-frequency differences that are more than five standard deviations away from the mean (leaving 1,023,667 SNP_s_), the test statistic drops to 1.08 and, hence, becomes insignificant. Therefore, except for a smattering of outliers, these standardized allele-frequency differences are close to normally distributed, and have a variance in line with our eigenvalue-based estimator of *F*_ST_.

**Figure 3.**
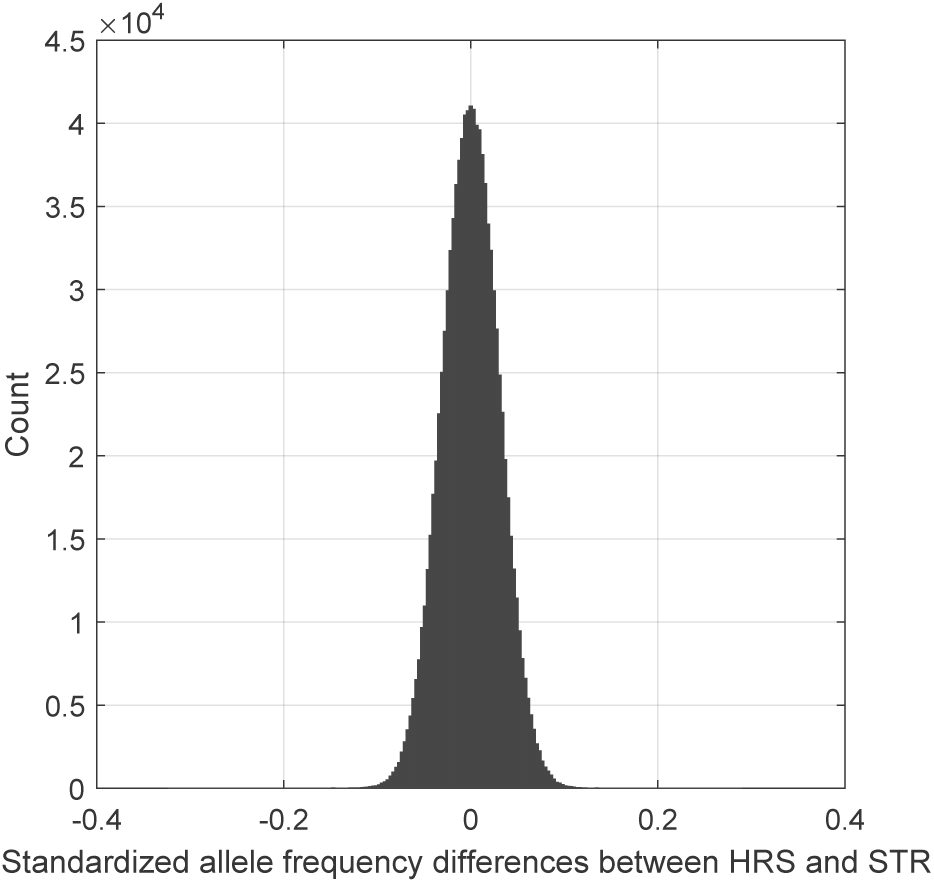
Histogram of standardized allele-frequency differences between the HRS and STR samples.

As least-squares techniques, such as LDSC regression, are sensitive to outliers – as an additional check – we inspect the average LD-score per LD-score percentile and plots these against the average GWAS *χ*^2^-test statistics, across SNP_s_ in these LD-score percentiles and across runs. Figure 4 shows the resulting scatter plot together with (I) simple-regression lines of the test statistics as explained by LD scores and (ii) lines predicted by theory, using the HRS-STR-specific estimate of *F*_ST_ (i.e., 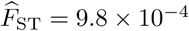). For the two lowest levels of stratification, we observe a strong agreement between predictions from theory and the fitted line based on average scores and statistics. However, for higher levels of stratification, we observe that the intercept is lower and the slope is higher than what the LDSC-regression theory predicts. In fact, according to theory, the slope should be independent from the amount of stratification. Yet, the slope of the fitted line under the highest amount of stratification is 74% higher than the slope under no stratification. This disparity lies reasonably close to the aforementioned inflation of ~ 84% in 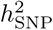 estimates from LDSC regression.

**Figure 4.**
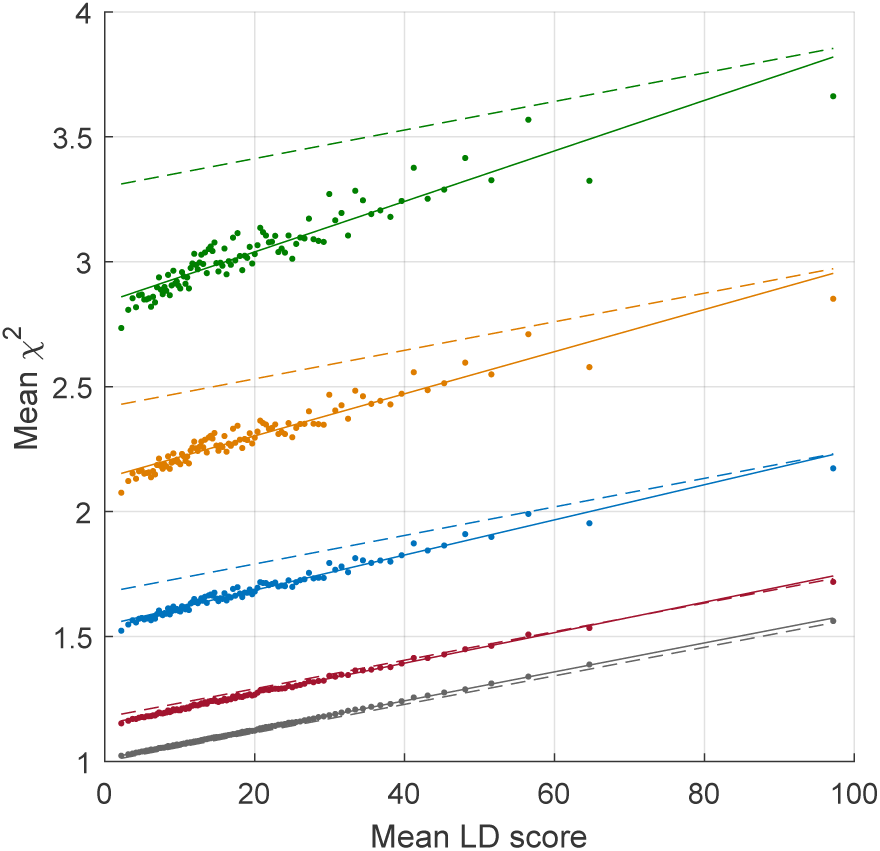
Scatter plot of the average LD score per LD-score percentile and the cross-run average *χ*^2^-test statistic per LD-score percentile. Gray dots: no stratification; red dots: light stratification; blue dots: moderate stratification; yellow dots: substantial stratification; green dots: strong stratification. Solid lines: fitted lines from a simple regression of mean test statistics on mean LD scores; dashed lines: predicted lines from LD-score-regression theory.

## Results for Three Discrete Populations

Figure 5 shows GREML and LDSC-regression estimates of the intercept and 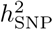 across the various runs and levels of stratification, for the simulations using three subpopulations. As with two populations, the intercept estimates are of the same scale and highly correlated (*R*^2^ = 99.15%).

**Figure 5.**
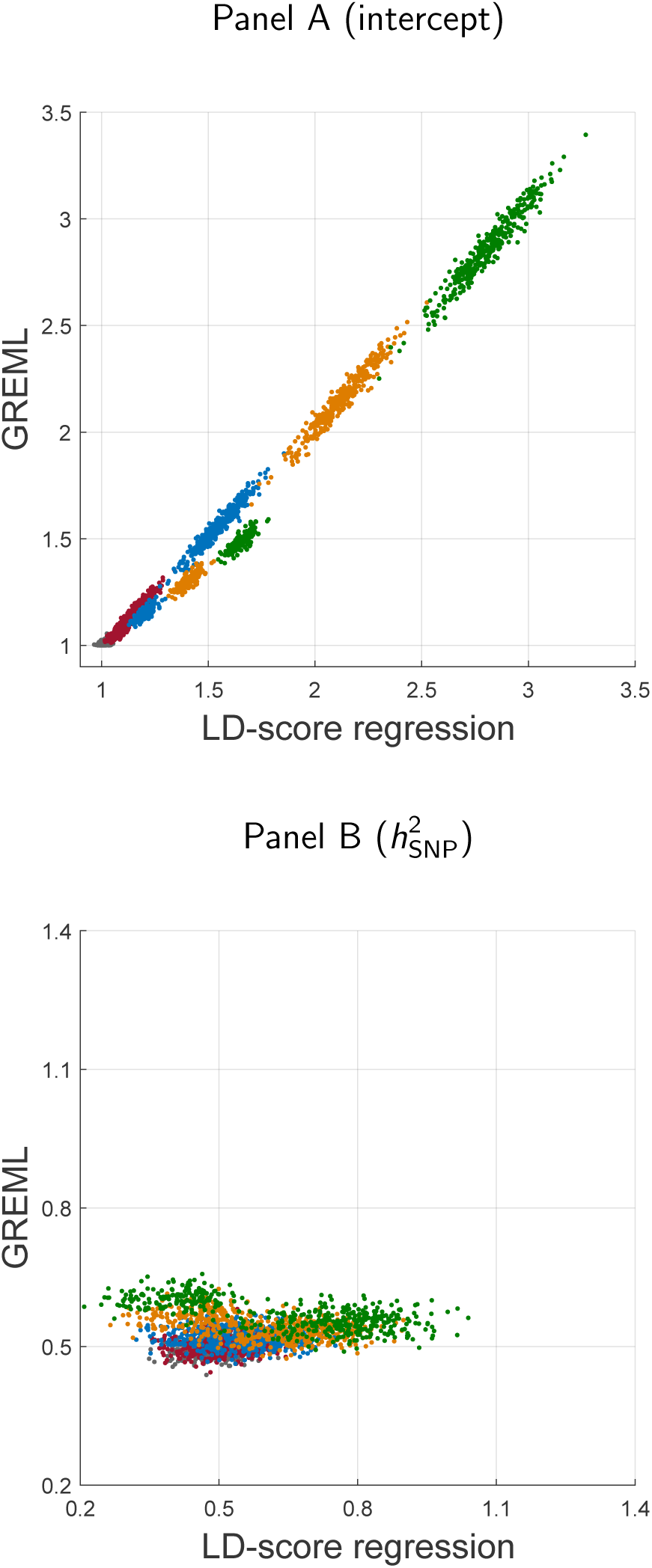
Scatter plots of estimates of the LD-score-regression intercept (Panel A) and SNP-based heritability (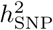; Panel B) across 500 runs and various levels of stratification based on data from *P* = 3 subpopulations. *x*-axis: LD-score regression estimates. *y*-axis: GREML estimates. In the GREML approach the two leading principal components are used as fixed-effect covariates, and the estimates of those effects 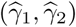 are cast to an intercept estimate using Equation 2. Gray dots: no stratification; red dots: light stratification; blue dots: moderate stratification; yellow dots: substantial stratification; green dots: strong stratification.

The reason why two slightly diverging clouds appear in the scatter plot is merely an artefact of the simulation. In the *P* = 2 case, one of the two subsamples gets assigned a positive mean whereas the other subsample gets assigned a negative mean (of the same magnitude as the positive mean). For comparability, in each run of the *P* = 3 case, each subsample gets assigned either the positive mean, the negative mean, or a zero mean. In runs where the two subsamples with the least genetic drift between them get assigned the negative and positive mean, this effectively leads to a smaller amount of stratification than intended *a priori* by simulation design. Nevertheless, under this scenario both LDSC and GREML intercept estimates are decreased, thereby, showing even more strongly that these two estimators are close to equivalent. It is important to point out though that in this particular case, GREML intercept estimates are on average a bit lower than the LDSC estimates. A likely explanation is that second PC becomes more important in estimating the intercept in this case. As Equation 2 is merely an extrapolation of Equation 1, the weight assigned to the squared coefficient for the second PC may therefore be underestimated.

The assertion of an underestimated weight for the second PC is supported by an additional regression of the LDSC-regression intercept estimates on 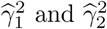 (i.e., the squared coefficients from the regressions of phenotypes on the leading two PCs). Figure 6 shows a scatter plot of the LDSC-regression estimates and a linear combination of 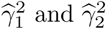, with weights based on this additional regression. The subtle divergence between estimates is now gone, and the *R*^2^ is up to 99.82%. Inspection of the weights reveals that Equation 2 indeed significantly underestimates the weight that ought to be given to the second PC; whereas Equation 2 sets the weight at 3.0 ×10^*−*4^, the optimal weight to retrieve the LDSC-regression intercept estimate as accurately as possible is 4.0 ×10^*−*4^ (s.e. = 2.7 ×10^*−*6^).

**Figure 6.**
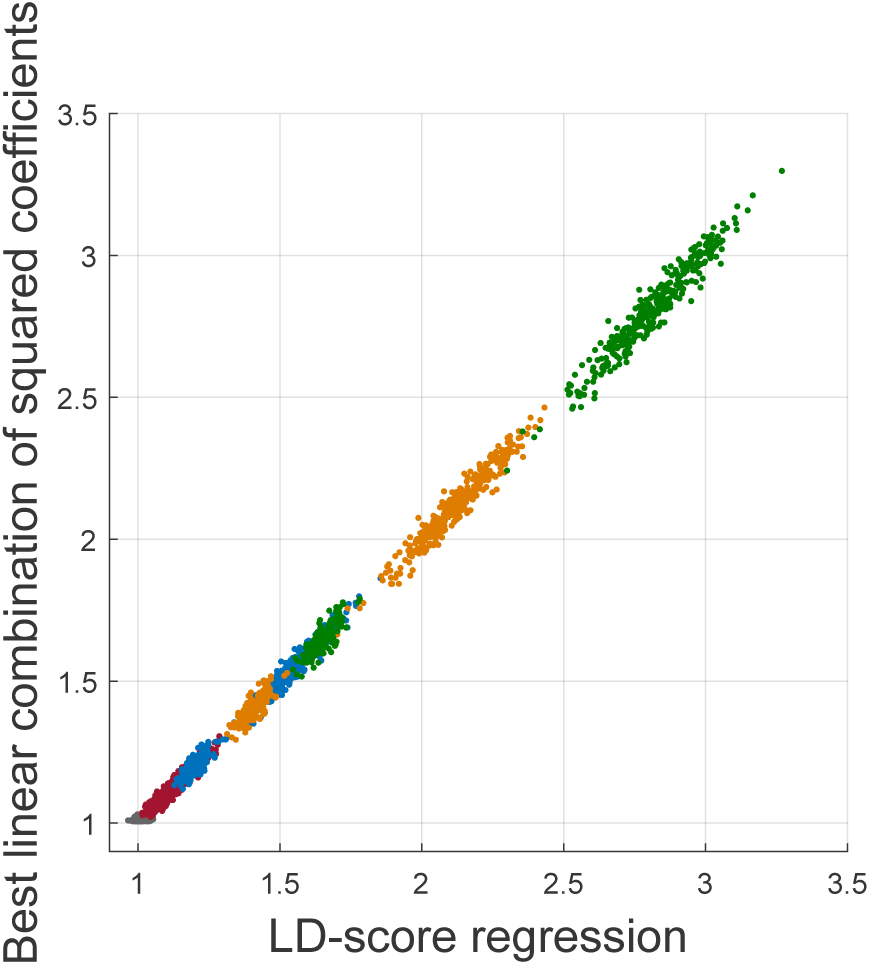
Scatter plot of estimates of the LD-score-regression intercept based on data from *P* = 3 subpopulations. *x*-axis: LD-score regression estimates. *y*-axis: a linear combination of squared coefficients of regressing the phenotypes on the leading two principal components and a vector of ones, with weights set by regressing the LDSC-intercept estimates on these squared coefficients and the vector of ones. Gray dots: no stratification; red dots: light stratification; blue dots: moderate stratification; yellow dots: substantial stratification; green dots: strong stratification.

Finally, we observe that the 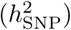 estimates from both methods are inflated with increasing amounts of stratification, and also that this bias seems to affect LDSC estimates more strongly than GREML estimates. Yet, this difference is less pronounced than what we observed in case *P* = 2. Including more PCs as control variables does not improve the situation; when including the five leading PCs of the GRM as fixed-effect control variables, under the highest level of stratification, the mean 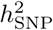 estimate from GREML estimation is still ~ 57%. Consequently, it seems that, even though in our simulations GREML 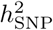 estimates are consistently less upwards biased than LDSC estimates, the additional bias of LDSC regression compared to GREML estimation is abated somewhat as the type of stratification becomes more complex.

## Empirical Results for Multiple Populations

We now consider two real phenotypes, *viz*., human height and body-mass index (BMI). Details on the QC are reported in Appendix B. As these phenotypes have been standardized at the study level (i.e., standardized to mean zero and unit variance in the STR, HRS, and RS samples separately before pooling data), the most important source of stratification has been eliminated. Consequently, we have clean traits for which we expect little stratification along the lead PCs. Hence, as the remaining stratification is likely to be of a higher order, we set *P* = 20. We study these phenotypes using LDSC regression and GREML estimation. In addition, we also perform semi-empirical analyses, in which we assign a phenotypic mean of +0.5 to the HRS samples, − 0.5 to the STR samples, and preserving the zero mean in the RS samples. In doing so, we introduce artificial stratification.

Table 3 shows estimates of the intercept and 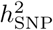 from both methods and both phenotypes, without additional stratification, and the two phenotypes, with added stratification. The intercept estimates from both methods are similar and – as expected – increase as stratification is added to the phenotypes. As sample sizes per subsample differ (e.g., for height we have 5,847 samples in HRS and only 4,328 in STR), these findings imply that Equations 1 and 2 are robust when sample sizes are not completely equal across subsamples. Furthermore, as the baseline phenotypes are real outcomes, subtle stratification may be at play (i.e., *P* may be large); if this is true, that would imply Equation 2 approximates the LD-score regression intercept quite well even when *P* is fairly large. In addition, the GREML 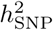 estimates seem credible; 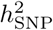 ~ 40% for height and ~ 19% for BMI. LDSC estimates are relatively similar. However, when introducing the additional stratification – in line with the simulation results for *P* = 2 – the LDSC 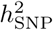 estimates increase considerably for both traits, whilst GREML 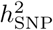 estimates remain relatively unperturbed. Hence, these findings provide further support of the notation that under strong stratification, some of the stratification may get absorbed by the slope of the *χ*^2^ statistics versus the LD scores.

**Table 3.** Intercept and SNP-based heritability 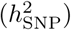 estimates from LD-score regression and GREML, using height and body-mass index (BMI), both standardized per subsample, and height and BMI, standardized per subsample and with artificial stratification added per subsample.

| Phenotype | Intercept | | $h_{\text{SNP}}^2$ | |
| --- | --- | --- | --- | --- |
|  | LDSC | GREML* | LDSC | GREML** |
| Height*** | 1.03 | 1.05 | 47.7% | 40.3% |
| Height + stratification**** | 2.26 | 2.35 | 65.7% | 41.1% |
| BMI*** | 1.02 | 1.01 | 12.5% | 19.4% |
| BMI + stratification**** | 2.02 | 2.08 | 35.1% | 21.2% |

\* Intercept estimate based on 20 leading PCs and eigenvalues using Equation 2
\*\* $h_{\text{SNP}}^2$ estimate obtained when controlling for 20 leading PCs
\*\*\* Standardized phenotypes per subsample
\*\*\*\* Phenotypic means adjusted to $-0.5$ in HRS and $+0.5$ in STR (RS: unchanged)

## Results for Different Sources of Stratification and Controls

Intercept and 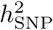 estimates from LDSC regression and GREML obtained using simulated phenotypes, under different sources of stratification and various controls for that stratification, are reported in Table 4. The data-generating process is identical to previous simulations for two discrete populations under strong stratification, except for one addition; here, we consider two sources of stratification, *viz*., (i) differences in phenotypic mean between the HRS and STR samples and (ii) differences in the phenotype shaped by the lead PC from the GRM. The first approach is, statistically speaking, equivalent to a sample dummy shaping phenotypic differences in mean.

**Table 4.** Average SNP-based heritability (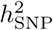) and intercept estimates across 500 runs and corresponding standard errors (s.e.), for GREML estimation and LDSC regression in HRS and STR data (i.e., *P* = 2) under strong stratification, shaped by either (i) sample dummies or (ii) the empirically inferred lead PC, and either controlled for by (i) the aformentioned dummies, (ii) the lead PC, or (iii) not controlled for at all.

| Method | Control* | $h_{\text{SNP}}^2$ estimates (s.e.) | | Intercept estimates (s.e.) | |
| --- | --- | --- | --- | --- | --- |
|  |  | Stratification shaped by sample dummy** | Stratification shaped by lead PC | Stratification shaped by sample dummy** | Stratification shaped by lead PC |
| LDSC | none | 94.28% ( 0.66% ) | 168.58% ( 0.74% ) | 2.825 ( 0.007 ) | 3.006 ( 0.007 ) |
| LDSC | sample dummy | 50.48% ( 0.35% ) | 63.81% ( 0.38% ) | 1.007 ( 0.001 ) | 1.073 ( 0.001 ) |
| LDSC | lead PC | 50.34% ( 0.35% ) | 50.50% ( 0.35% ) | 1.035 ( 0.001 ) | 1.006 ( 0.001 ) |
| GREML | sample dummy | 50.29% ( 0.13% ) | 58.59% ( 0.12% ) | 3.293 ( 0.006 ) | 1.554 ( 0.004 ) |
| GREML | lead PC | 53.72% ( 0.13% ) | 50.28% ( 0.13% ) | 2.918 ( 0.008 ) | 3.293 ( 0.008 ) |

\* Control for LDSC: covariate in the GWAS prior to LDSC regression; control for GREML: fixed-effect covariate in GREML estimation
\*\* Stratification shaped by sample dummies (i.e., HRS and STR dummies) is equivalent to assigning differences in mean between samples

In terms of controls, we apply LDSC regression to three different sets of GWAS results, *viz*., (i) with no controls in the GWAS, to further assert whether LDSC regression is able to deal with different forms of population stratification, (ii) controlling for the aforementioned sample dummy, (iii) controlling for the lead PC. For GREML estimation, we use (i) the sample dummy (scaled to unit length and mean zero) to control for stratification and estimate the intercept, and (ii) the lead PC to control for stratification and estimate the intercept.

Importantly, the squared correlation of the sample dummy and the lead PC is 83.75%. This high correlation implies that, in cases where the control for stratification – provided a control is present – differs from the source, only some residual stratification is left. For GREML, the inflation in 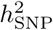 estimates is symmetric when there is residual stratification. That is, when either the sample dummy shapes stratification and the lead PC is used as control or *vice versa*, 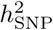 estimates are biased upwards significantly.

For LDSC regression the results are more complex. First, we observe – as before – that LDSC regression yields highly inflated 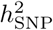 estimates when there is strong stratification in the GWAS results (i.e., when the stratification is not controlled for in the GWAS stage), regardless of whether that stratification is shaped by the sample dummy or the lead PC. However, when we attempt to correct for the stratification in the GWAS by an imperfect control, an asymmetry arises. More specifically, when the stratification is shaped by the empirically inferred lead PC, but controlled for using the sample dummy, LDSC regression is not fully able to accommodate the residual stratification; the average 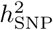 estimate across 500 runs of simulations then equals ~ 63.8% (s.e. = 0.4%). On the other hand, when the stratification is shaped by differences in mean between the HRS and STR samples (i.e., the sample dummy), yet is controlled for in the GWAS using the lead PC, LDSC regression seems able to address the residual stratification; the average 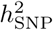 estimate across 500 runs then equals ~ 50.3% (s.e. = 0.4%).

From a theoretical perspective this asymmetry is not surprising; LDSC regression assumes two discrete populations with differences in mean (i.e., a population dummy shaping the stratification) and the lead PC just tends to correlate highly with such a dummy variable. Therefore, when the stratification is shaped by such a dummy, the lead PC will take out most of the stratification, yet some stratification will be left. However, that residual is still shaped by the population dummy—the type of stratification that LDSC regression is built to address. Hence, LDSC regression picks up that residuals and effectively ‘controls’ for it. Conversely, when stratification is shaped by the lead PC, but controlled for using the population dummy in the GWAS, the residual is effectively outside the scope of what LDSC regression can deal with, thereby, inflating 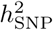 estimates even when that residual is relatively small.

## Interpretation of the LD-Score-Regression Intercept Estimated Using Individual-Level Data

Based on the broad set of simulations and empirical analyses, we conclude that the LDSC-regression intercept can be approximated with high precision by a weighted sum of squared regression coefficient of the standardized phenotype on the leading PCs from the GRM. These weights increase with the corresponding eigenvalues, which in turn increase with the amount of genetic drift. The squared regression coefficients from the PCs can be written as the product of sample size and the *R*^2^ of that PC with respect to the phenotype. Although a regression of the LDSC estimates on these squared coefficients reveals that the weights given to these squared coefficients, in Equation 2, are slightly off-target for higher-order stratification (i.e., not residing in the leading PC), the approximation is still fairly accurate. This assertion is corroborated by our empirical results.

These finding indicate that in individual-level data, the intercept is simply an increasing function of (i) the amount of drift in allele frequencies across subpoulations, (ii) the proportion of phenotypic variance explained by stratification, as tagged by the leading PCs, and (iii) sample size. These aspects are all in line with what one can expect intuitively based on the LDSC-regression derivations, where the intercept is also a linearly increasing function of sample size, genetic drift as shaped by *F*_ST_, and the squared difference in phenotypic mean across two subpopulations.

## Implications of Inflated SNP-Based Heritability Estimates

This study follows the data-generating process assumed in the original LDSC-regression derivations closely; we simulate standardized phenotypes using an infinitesimal model with standardized SNP_s_ having homoskedastic effects, with fixed cross-population differences in phenotypic mean, and drift in line with assumptions of LDSC regression^4^. Despite these efforts, under considerable stratification, we observe an intercept below expectation and a slope above expectation. There is no dimension along which our simulations strongly differ from the assumptions in the derivations of LDSC regression^4^. Consequently, LDSC estimates exhibit unexpected properties in extreme scenarios. More specifically, 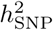 is significantly overestimated even under an intercept estimate as low as 1.122 (Table 2; under *P* = 3 and light stratification). Importantly, the corresponding attenuation ratio^19^, defined as the estimated intercept minus one and the average *χ*^2^-test statistic minus one, has a mean of 0.388 across runs, indicating that ~ 39% of the inflation in test statistics can be attributed to stratification in this simulation design. This result implies that, even under our ‘light stratification’ design in three discrete samples, the GWAS test statistics are already distorted considerably by stratification. Hence, as much of the inflation in test statistics can be attributed to stratification, it is not entirely surprising that estimation of 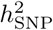 goes awry. However, in spite of this considerable ratio, our findings do imply that LDSC-regression estimates should be interpreted with caution, at the very least, when the intercept is significantly different from one.

Although we consider substantial differences in phenotypic mean and, therefore, intercepts significantly larger than one, a more reasonable scenario may be conceived where similar dynamics play a role, *viz*., in very large samples, where even a subtle difference in phenotypic mean across subpopulations can result in a large intercept, as the intercept is an increasing function of sample size. Our results suggest that under such a scenario, the intercept may get underestimated and the slope overestimated, inflating 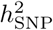 estimates. Hence, further research on the asymptotic properties of LDSC regression is warranted.

In light of our findings, we agree with the assertion by Bulik-Sullivan et al. (2015) that “*whenever possible, it is preferable to obtain all relevant genotype data and correct for confounding biases directly; post-hoc correction of test statistics is no substitute for diligent quality control*”^4^. LDSC regression is not a panacea for population stratification; it can only deal with a narrowly defined and limited amount of confounding stratification. However, provided population stratification is carefully controlled for in the GWAS stage, LDSC regression remains an informative tool for inferring the amount of residual stratification permeating GWAS summary statistics *post hoc*.

## Appendices

### A Derivations Estimator for Individual-Level Data

We first recapitulate the stratification assumed in the derivations of LDSC regression^4^ and generalize to *P* discrete populations. Based thereon, we derive an unconditional expected GRM. By assuming that the average magnitude of the drift away from pooled allele frequencies is the same across the populations (which holds by definition in the two-populations-based theory underpinning LDSC regression), we can derive a closed-form expression of the eigendecomposition of the expected GRM. We show that all eigenvalues – except the leading *P* - 1 eigenvalues – are decreased by the same small amount due to stratification, whereas, the leading *P* - 1 eigenvalues are increasing functions of sample size as a result of drift. For *P* = 2, we derive an explicit transformation of the estimated association between the first PC and the phenotype, providing an estimate of the LD-score regression intercept.

#### A.1 Genetic Drift in LD-Score Regression

In the derivations of LDSC regression^4^, stratification is conceptualized as a GWAS sample consisting of individuals drawn from two independent populations, with different allele frequencies due to drift and different phenotypic means. Using modified notation, the following is assumed:

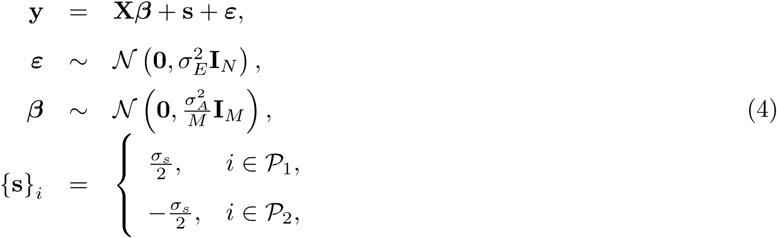

where 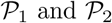 denote the sets of individuals drawn from Populations 1 and 2 respectively, where 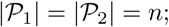; there are *n* individuals drawn from both populations, yielding *N* = 2*n* observations in total. Parameter *a*, found in Equation 2.14 of the Supplementary Note by Bulik-Sullivan et al. (2015)^4^, is approximately equal to 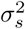 (i.e., the squared difference in the phenotypic mean of the subpopulations). **X** denotes an *N × M* matrix of *M* standardized SNP_s_, with random effects in vector ***β***. In line with preceding work^9^, another assumption is that effects of standardized SNP_s_, ***β***, are independent draws from a normal homoskedastic distribution. In addition, it is assumed that the effects in ***β*** are constant across the two populations. Vector ***ε*** denotes the environmental effects. Finally, 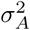 denotes the additive genetic variance and 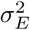 the environment variance, and consequently 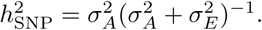.

Let *f*_1_ and *f*_2_ denote the allele frequency for a given SNP in Populations 1 and 2, and 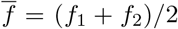 the average allele frequency across the two populations. For now, assume that all distributions, expectations, and variances, are conditional on *f*_1_, *f*_2_, and, thereby, on 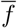. The additively-coded genotype for individual *i* (denoted by *g*_*i*_ ∈ {0, 1, 2}) then satisfies the following properties

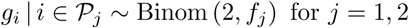

Therefore,

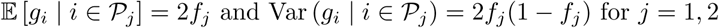

Moreover, as indicated, individuals are sampled from the two populations with equal chance. Hence, *g*_*i*_ is a draw from a mixture distribution with mean 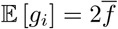 and variance

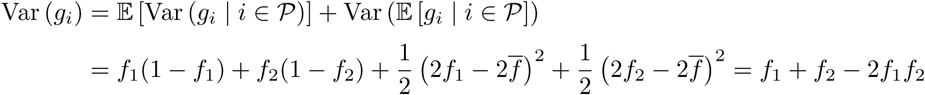

Genotypes are standardized, such that the standardized genotype of individual *i* (denoted by *x_i_*) is given by

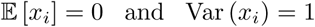

Therefore, LDSC regression^4^ implicitly assumes that

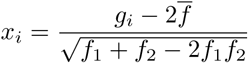

where 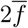 and *f*_1_ + *f*_2_ − 2*f*_1_*f*_2_ are the theoretical expectation and variance respectively, of a random variable that is drawn from the aforementioned mixture distribution. In this case, we have that

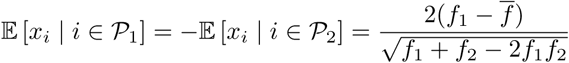

Conditioning on the pooled frequency 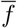, LDSC regression implicitly assumes

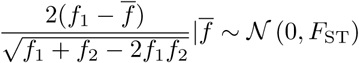

where *F*_ST_ denotes Wright's *F*-statistic^15^, measuring the amount of genetic drift across populations. The larger *F*_ST_ the larger, on average, the allele frequency differences across populations will be. Provided allele frequencies do not vary too much across the populations, 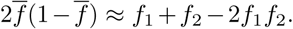 For instance, when the cross-population difference in allele frequency is 10% and the pooled frequency is 50%, we have that 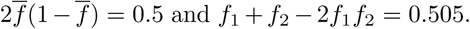 Thus, even at a substantial allele frequency difference, the relative difference between the true variance, given by *f*_1_ + *f*_2_ − 2*f*_1_*f*_2_, and the variance inferred by 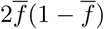 is only 1%. Therefore, we make a slight amendment, and assume that

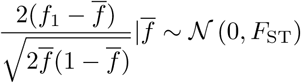

This change greatly simplifies the closed-form solution of the GRM. More precisely, the GRM (e.g., constructed using GCTA^9^) is constructed assuming each SNP is binomially distributed (i.e., in Hardy-Weinberg equilibrium; HWE). Therefore, each SNP is standardized according to its pooled – potentially admixed – allele frequency 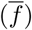 assuming HWE (i.e., in addition to correcting for the expected value of the raw genotype, given by 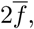, the raw genotypes are also divided by 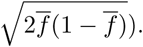 By replacing *f*_1_ + *f*_2_ − 2*f*_1_*f*_2_ by 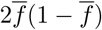 in the preceding expression, when deriving the unconditional expectation of the GRM, the denominator in the left-hand-side of this expression and the standardizing coefficient of the SNP, when constructing the GRM, cancel each other out.

The distribution of the difference in allele frequency between the first population and the pooled frequency, can now be written as

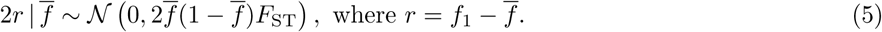

Hence, 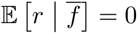 and

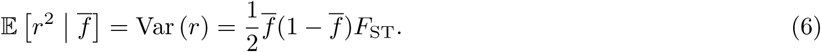

This expected squared difference between the population-specific and the pooled allele-frequency is in line the updated Nei estimator of *F*_ST_^16,21^. In the derivation of LDSC regression, when discussing the distribution of the standardized difference in allele frequency, the same literature is pointed to. Moreover, the preceding expression aligns with an expression that is referred to as the “*most common explicit computational formula*” for *F*_ST_^22^.

This expression explicitly accounts for the loss of one degree of freedom across populations, when considering deviations from the pooled allele frequency. We should point here that in later work the loss of this degree of freedom is ignored^23^, which would correspond to

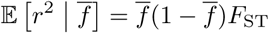

This approach is adopted – for instance – in related work^24^. Rather than commenting on whether one should account for the lost degree of freedom or not, our focus should be to keep as close as possible to the LDSC-regression approach^4^. Therefore, we assume that the variance of *r* is as given in Equation 6. Although we make the implicit distribution of 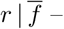 assumed in the derivations of LDSC regression^4^ – explicit, without loss of comparability of our methods, further derivations show that we need to make no assumptions about the type distribution of 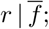 the only thing we need to impose is the condition that its expectation is zero and the variance as shown in Equation 6.

#### A.2 Drift and Stratification in More than Two Populations

Letting **f** denote a *P ×* 1 vector of allele frequencies in *P* populations, Equation 5 can be generalized as

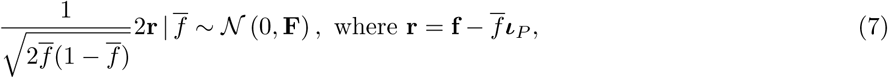

where ***ι**_P_* is *P*-dimensional column vector of ones, and where **F** is *P × P* matrix of *F*-statistics, for which the diagonal elements indicate the amount of drift away from 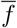 for each population and the off-diagonal elements indicate the extent to which different populations covary in their drift away from 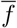. In LDSC regression we have *P* = 2 and

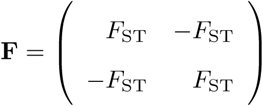

Without loss of generality, we can order the phenotype vector, **y**, according to the populations from which the individuals are drawn. As indicated, we consider SNP_s_ that are standardized according to cross-population allele frequencies. That is, for individual *i* and SNP *k*, the standardized genotype is given by

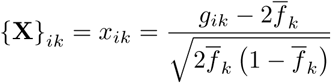

where *g*_*ip*_ ∈ 0, 1, 2} is the additively-coded genotype. Generalizing Equation 4 to *P* populations, and rewriting it in terms of variance components, the GRM, and a vector of phenotypic means per populations, we have:

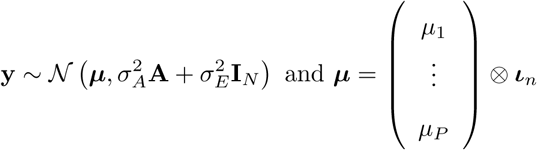

where **A** = *M* ^−1^**XX**^*T*^ denotes the *N × N* GRM in the admixed sample, estimated from *M* markers, and **I**_*N*_ the identity matrix of appropriate dimensions, where *N* = *nP*. In the original LD-score regression framework, we have that *µ*_1_ = 2^*−*1^*σ_s_* and *µ*_2_ = − 2^*−*1^*σ_s_*.

#### A.3 Expected GRM and Eigendecomposition under Drift

We now consider the expectation of the GRM. We first derive the elements of the expected GRM based on a single SNP, conditional on the pooled and within-population allele frequencies (i.e., 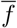, *f*_1_, and *f*_2_), for *n* = 2, and omitting subscripts for the index of the SNP. By applying the law of iterated expectations to each element, we obtain the expected GRM independent of allele frequencies. Using this allele-frequency-independent expected GRM, we can generalize to an *N*-by-*N* GRM with *n >* 2 individuals from *P* populations.

As indicated, each SNP is standardized under the assumption of HWE. That is, the standardized genotype of individual *i* for a given SNP with pooled allele frequency 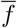, is given by

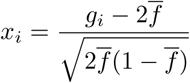

First note that, for 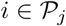

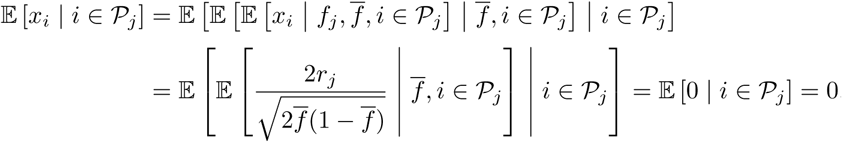

where *r*_*j*_ is the *j*-th element of **r**. Now, the expected relatedness of individual 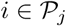 with itself, is given by

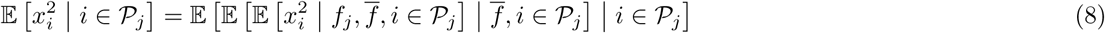

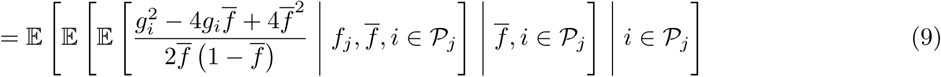

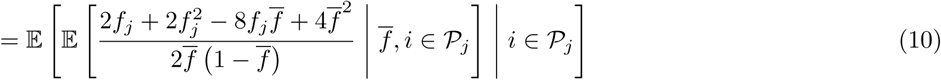

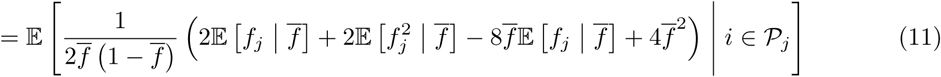

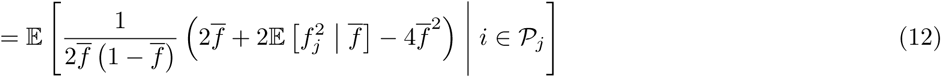

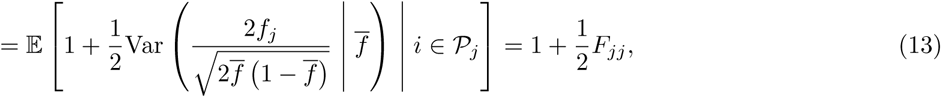

where *F_jh_* is element {j, h} of **F**. Similarly, for individuals *i* ≠ *l* from Populations *j* and *h*, we have

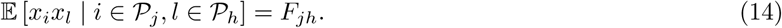

Hence, the expected GRM with *n* observations per population, sorting individuals by population index, is

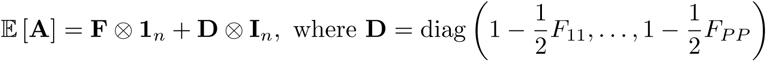

and where **1**_*n*_ is *n × n* matrix of ones and ⊗ denotes the Kronecker product.

Assuming *F*_*jj*_ = *F*_0_ ≥ 0 for *j* = 1, …, *P* (i.e., the magnitude of drift away from the pooled frequency is equal across populations; an assumption which holds by definition when *P* = 2), the diagonal elements of **F** are equal. Under this assumption, we have

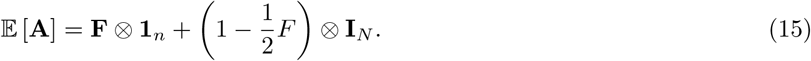

Letting the eigendecompositions of the matrix of *F*-statistics and ones be given by

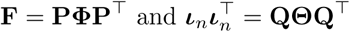

then

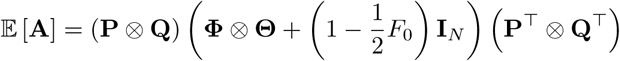

By construction 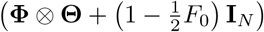 is a diagonal matrix and (**P** ⊗ **Q**) is an orthonormal matrix. Hence, 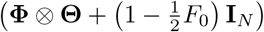 is a diagonal matrix containing the eigenvalues of 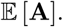 Inspection of 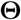 reveals that

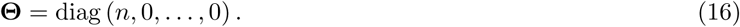

As **F** is a covariance matrix its eigenvalues are non-negative. The eigenvalues of the expected GRM, in descending order, are now given by

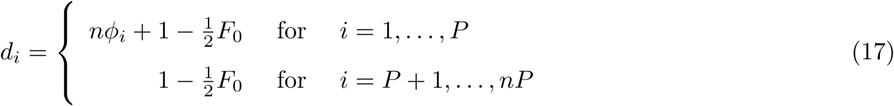

In case *P* is small and 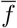 is the empirical midpoint of frequencies in the two populations, matrix **F** is unlikely to have full rank. For instance, in case *P* = 2, we have that *F*_11_ = *F*_22_ = *F*_0_ = *F*_ST_ and *F*_12_ = −F_ST_, in which case *φ*_1_ = 2*F*_ST_ and *φ*_2_ = 0, and thereby 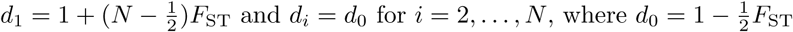 and *N* = 2*n*.

Focussing on the case where *P* = 2, the first eigenvalue of the expected GRM is affected by the product of the total sample size and Wright's *F*-statistics. Even for fairly small values of *F* _ST_, this quantity grows large with increasing sample sizes. The remaining eigenvalues, however, are not affected by sample size; each remaining eigenvalue is merely decreased by 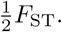 Values of *F*_ST_ are usually small (e.g., it is suggested that *F*_ST_ ≈ 0.01 for populations on the same continent^4^). Under this approximation, all remaining eigenvalues would be approximately equal to one.

#### A.4 Least-Squares-Based Estimator of the Intercept

The log-likelihood function of GREML estimation^9^, ignoring the constant and including the leading PCs as fixed-effect covariates, is given by

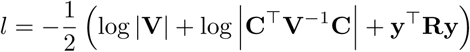

where **C** is the matrix of fixed-effects covariates, **V** is the phenotypic covariance matrix, and where

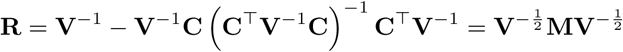

where **M** is an idempotent matrix, projecting onto the null space of 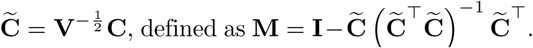

In our case **C** is merely a vector, defined as the first PC from the GRM. Hence, we switch to lower-case notation c, and replace c by its theoretical expression, which is given by the Kronecker product of the first column of **P** and of **Q**. That is,

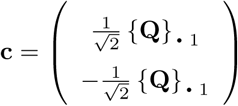

where 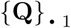 denotes the first column of **Q.** Bearing in mind that matrix 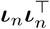 has rank one, its first eigenvalue and eigenvector are sufficient for reconstructing 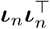 This observations implies that

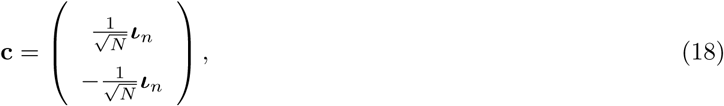

Rewriting the original model in Equation 4, we have

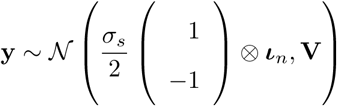

where 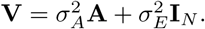 Replacing **A** by its expectation under stratification, **V** can be rewritten as

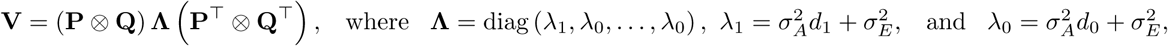

with *d*_1_ and *d*_0_ as defined in Appendix A, where the *d*_1_ increases with sample size as a result of drift, while *d*_0_ ≈ 1 provided *F*_ST_ is small. Now 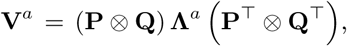 where the Λ^a^ can be obtained by raising the diagonal entries of Λ to the power a. Hence 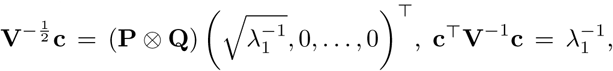 and 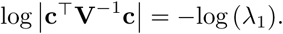 Based on these expressions, we can show that

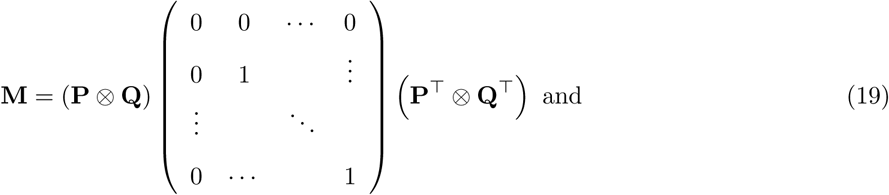

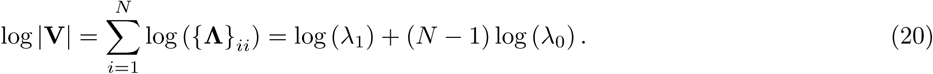

Consequently, we can now write the log-likelihood as follows

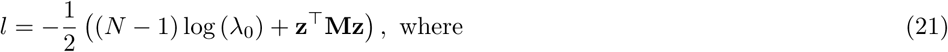

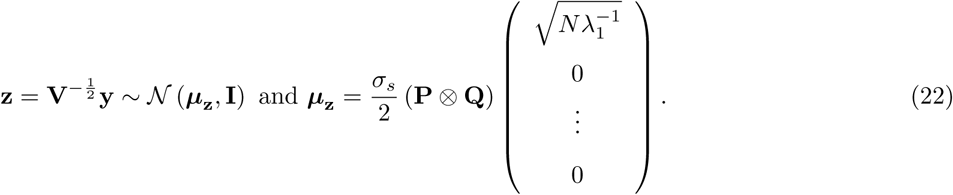

Exploiting the fact that **M** is idempotent, we have that

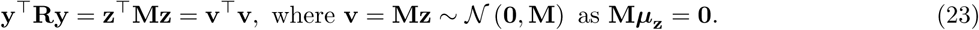

Hence, by including the first PC as fixed-effect covariate, the population-dependent mean is eliminated from the likelihood. Moreover, the term 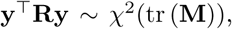, where tr (**M**) = *N* − 1. By rewriting **M** in terms of individual PCs, rather than a Kronecker product, we can show that 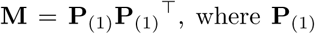 denotes the matrix of all eigenvectors from the expected GRM except the first. In the last expression for the log-likelihood, the leading eigenvalue has also been eliminated from the combined term log 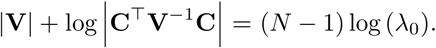 Consequently, the GREML log-likelihood obtained by including the first PC from the GRM as fixed-effect covariate, is independent of the first eigenvalue and first eigenvector of the GRM. Since the effect of stratification on the first eigenvalue is of the order *NF*_ST_, whilst the effect on other eigenvalues is only of the order *F*_ST_, it is obvious that this approach will remove the vast majority of any potential bias incurred due to stratification. Hence, we posit that – under the same data-generating process assumed in the derivation of LDSC regression – GREML estimation including the first PC as fixed-effect covariate will be approximately unbiased, provided *F*_ST_ is small.

We will now study the expected value of the fixed-effect estimate of the first PC using generalized least squares (GLS) and ordinary least squares (OLS), and study its relation with the LD-score regression intercept. The GLS (or REML fixed-effects) estimator is given by

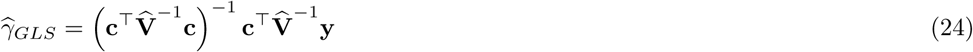

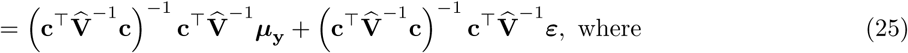

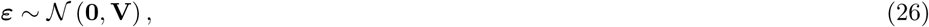

where 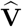 denotes the estimate of the true covariance matrix **V**, based on estimates 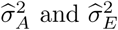 of the true variance components 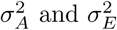 of the model. Now,

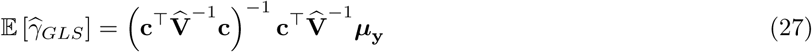

Substituting expressions found before, we have that

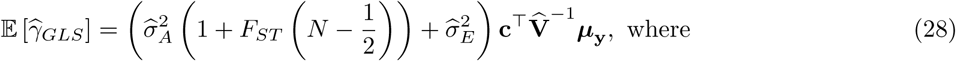

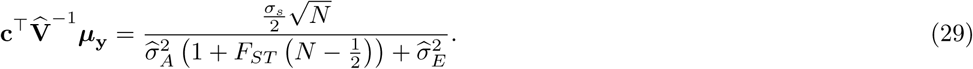

Therefore,

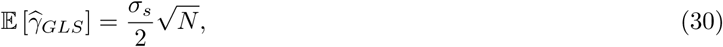

where *N* = 2*n* denotes the total sample size. Owing to the unit length of the PCs, the OLS estimator is given by 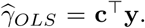 We show in Appendix D, that, in this particular model, 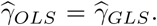 Hence, regressing the phenotype on the first PC using OLS is just as efficient as using GLS in this particular instance. Hence, we omit the OLS and GLS subscript from this point on. Using the fact that 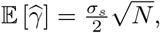 we have that

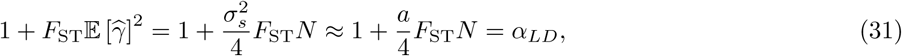

where 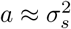 is the squared difference in phenotypic mean between the two subpopulations, and where *α*_*LD*_ denotes the theoretical LD-score-regression intercept, given *a*, *F*_ST_, and *N*. This theoretical expression is based directly on Equation 2.14 in Section 2 of the Supplementary Note to the LDSC-regression derivations^4^, taking into account an error in Equation 2.11 (in which the right-hand side should be equal to 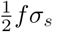) and the knock-on effects of a correction for this mistake. Using the fact that the first eigenvalue of the GRM is loosely expected to be given by

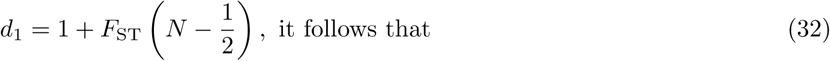

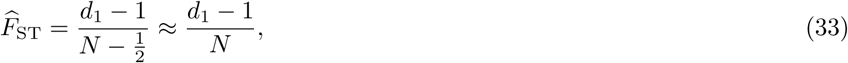

where the (notationally neater) approximation of *N* − 0.5 by *N* hardly affects the estimate of 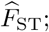 when *N* is as low as 100, the approximation on the right-hand side is only 0.5% lower than the initial expression. When, more realistically, *N* > 10k, the approximation of 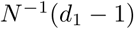 is only 0.005% lower than (*N* − 0.5)^*−*1^(*d*_1_ − 1).

Combining terms, our individual-level-data-based estimator of the LDSC-regression intercept is given by

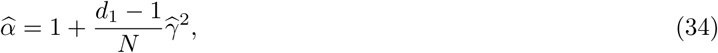

where *d*_1_ denotes the first eigenvalue from the GRM, 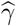 the estimate of the regression of the phenotype on the first PC from the GRM, and *N* the total sample size. Importantly, we tacitly assume the phenotype to be standardized to have mean zero and unit variance. Of course, 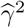 is not an unbiased estimator of 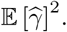 However, an unbiased estimate of 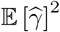 hinges on knowing the true variance components. Nevertheless, we would like to point out that

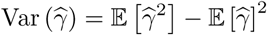

Hence, in our approximation, where the squared expectation of the estimator is replaced by the squared estimate, this squared estimate has the following expectation

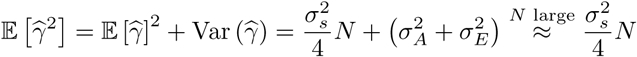

where, for derivational ease, 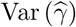 is set equal 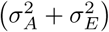 (i.e., the phenotypic variance after subtracting the variance accounted for by the leading PC). As the phenotype is standardized, 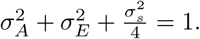 Hence, 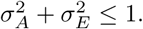 Consequently, provided the sample is sufficiently large, 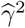 is an acceptable estimator of 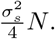

Writing the linear mixed model as follows

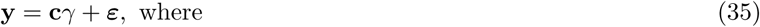

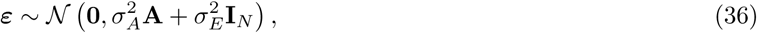

where **c** denotes the leading principal components, we can see both the OLS and GLS estimator aim to estimate *γ*, and from this draw inferences about the LD-score regression intercept. OLS ignores the structure of the covariance matrix of ***ε***, where GLS assumes *σ*_*A*_ and 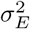 – or estimates thereof – are given. As it is possible to write down a linear regression function for the expectation of pairwise phenotypic products between individuals (i.e., *y*_*i*_*y*_*j*_ for *i* = 1, …, *N* and *j* = 1, …, *N*), in this mixed model with one fixed-effect regressor, an extended version of a Haseman-Elston regression can also be applied; the extension here being that pairwise products of loadings on the first PC between individuals then need to be included as regressor (i.e., *c*_*i*_*c*_*j*_). The estimated effect of *c*_*i*_*c*_*j*_, also denoted by 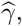 can then be cast to an intercept estimate using Equation 34.

### B Quality Control

The pooled genotype data, which is used as basis for the simulation study (discussed in Appendix C), is described in detail in the Supporting Information of existing work^17^. Summarizing, the Swedish Twin Registry (STR) samples have been genotyped using the HumanOmniExpress 12 v1 array, the Health and Retirement Study (HRS) samples using the Infinium Omni 2.5 array, and the Rotterdam Study (RS) samples using Illumina HumanHap 550K array. The STR and RS samples have been imputed `MaCH/Minimac` and HRS samples using `IMPUTE2`. Hence, there are both differences in the genotyping array and in the imputation procedure. The three studies have been imputed using the 1000 Genomes reference panel^25^.

Regarding quality control (QC) prior to creating this pooled dataset, in the imputed samples only HapMap 3 SNP_s_^18^ are selected. Genotypes are hard-called by rounding the dosages. Only high-quality SNP_s_ are selected (e.g., low missingness, high imputation quality). For full details see the first QC stage in Supporting Information, S1 Data, of the work by De Vlaming et al. (2016)^17^. For the purposes of our simulation we apply additional QC. The pooled dataset comprises 8,652 individuals of North-West European ancestry from the HRS and an additional 9,617 individuals from the STR. There are 1,062,589 SNP_s_ prior to additional QC.

The additional QC steps we apply are as follows:

1. We exclude the 26 regions reported in Table 5. This list is based on regions known to harbor inversions^26^ and is supplemented with additional regions found using InvFEST^27^. Excluding such large inversions is important, as these regions induce long-range LD and would, therefore, strongly affect leading principal components from genetic data if not removed.
2. We exclude SNP_s_ with
  - any missingness,
  - a minor allele frequency below 1%, and/or
  - a Hardy-Weinberg-Equilibrium-test *p*-value below 10^−6^.
3. We apply a relatedness cut-off of 0.025 using `PLINK`.
4. As the three studies differ in sample size, we select the largest possible random subsample per study, such that the sample size is equal for each of the three subsamples in the pooled data.
5. We again exclude SNP_s_ with a minor allele frequency below 1%, and/or a Hardy-Weinberg-Equilibrium-test *p*-value below 10^*−*6^.
6. We exclude SNP_s_ that are not available for the European-ancestry samples in the 1000Genomes, Phase 3 reference panel^28^, as available at https://data.broadinstitute.org/alkesgroup/LDSCORE/1000G_Phase3_plinkfiles.tgz (accessed on July 26, 2017).
7. For the empirical analyses only: we exclude individuals for whom the phenotype of interest is not available.

**Table 5.** Regions in the human genome excluded, with basepair positions according human-genome build 37.

| Chromosome | Basepair Position |  |
| --- | --- | --- |
|  | Start | End |
| 1 | 48,287,980 | 52,287,979 |
| 2 | 86,088,342 | 101,041,482 |
| 2 | 134,666,268 | 138,166,268 |
| 2 | 183,174,494 | 190,174,494 |
| 3 | 47,524,996 | 50,024,996 |
| 3 | 83,417,310 | 96,017,310 |
| 5 | 44,464,243 | 50,464,243 |
| 5 | 97,972,100 | 100,472,101 |
| 5 | 128,972,101 | 131,972,101 |
| 5 | 135,472,101 | 138,472,101 |
| 6 | 25,392,021 | 33,392,022 |
| 6 | 56,892,041 | 63,942,041 |
| 6 | 139,958,307 | 142,458,307 |
| 7 | 55,225,791 | 66,555,850 |
| 8 | 7,962,590 | 11,962,591 |
| 8 | 42,880,843 | 49,837,447 |
| 8 | 111,930,824 | 114,930,824 |
| 10 | 36,959,994 | 43,679,994 |
| 11 | 46,043,424 | 57,243,424 |
| 11 | 87,860,352 | 90,860,352 |
| 12 | 33,108,733 | 41,713,733 |
| 12 | 111,037,280 | 113,537,280 |
| 17 | 31,799,963 | 33,389,579 |
| 17 | 40,928,985 | 42,139,672 |
| 20 | 32,536,339 | 35,066,586 |

After QC we have 17,544 observations for our simulation study, of which 5,848 from each of the three underlying studies (i.e., the STR, HRS, and RS). For each observation we have 1,023,716 SNP_s_ meeting our QC criteria.

In the empirical exercise we consider human height and body-mass index (BMI). For details on the construction and QC of these phenotypes we refer to earlier work^17^. Summarizing, these phenotypes have been aggregated across available measurements (when available), corrected for non-linear birth-year effects and sex, standardized to have mean zero and unit variance in each of the three subsamples separately, and pooled across studies thereafter, yielding *N* = 26, 448 for height and *N* = 26, 438 for BMI. After these QC steps, the phenotypes are merged with genetic dataset, leaving *N* = 15, 966 for height and *N* = 15, 959 observations for BMI. No further standardization is applied after merging the phenotype and genotype data. In these empirical analyses, we have unequal sample sizes per study per phenotype. Final sample sizes per phenotype per study are reported in Table 6. As the sample sizes differ quite substantially, the empirical exercise tests indirectly whether a violation of the equal-sample-size per subpopulations affects the relation between the LD-score-regression intercept and our individual-level-data estimator.

**Table 6.** Sample size for phenotypes height and body-mass index (BMI) per study after QC.

| Study | Phenotype |  |
| --- | --- | --- |
|  | Height | BMI |
| HRS | 5,847 | 5,845 |
| RS | 5,737 | 5,732 |
| STR | 4,382 | 4,382 |
| Total | 15,966 | 15,959 |

For the construction of LD-scores, we used the binary PLINK files available at the website of LD-score regression^4^, for the European-ancestry samples in the 1000Genomes, Phase III reference panel^28^, as available at https://data.broadinstitute.org/alkesgroup/LDSCORE/1000G_Phase3_plinkfiles.tgz (accessed on July 26, 2017).

LD-scores are constructed per chromosome with a one-centimorgan window using LD-score regression^4^, based only on the subset of SNP_s_ available at the end of the QC procedure for the pooled HRS-STR-RS dataset.

### C Simulation Study and Empirical Analyses

#### C.1 Two Populations

We use the pooled imputed genotype data from the HRS and STR, obtained after the quality control procedure, discussed in Appendix B. Based on this dataset we simulate phenotypes by means of an infinitesimal model, for various degrees of population stratification. More specifically, let *r* = 1, …, 500 denote the index of the runs, *α* = {0, 0.25, 0.5, 0.75, 1} the set of stratification levels (index by *l* = 1, …, 5), and 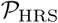 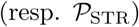 the set of HRS (STR) individuals. We simulate the phenotype for individual *i* in run *r* for stratification level *l* as follows:

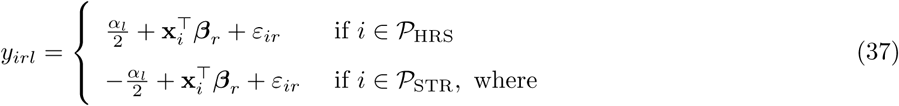

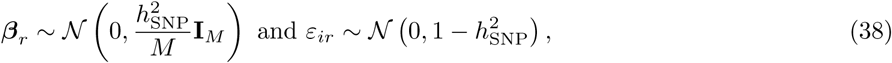

where *α_l_* denotes the *l*-th element of set *α* and 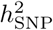 denotes the SNP-based heritability. All random draws in vectors ***β**_r_* and scalars *ε*_*ir*_ are independent from each other. Scalar *M* denotes the number of SNP_s_, and **x**_*i*_ denotes the *M ×* 1 vector of genotypes for individual *i*, standardized at the level of the pooled sample (assuming Hardy-Weinberg equilibrium holds). The standardized genotype of individual *i* for SNP *k* (denoted by *x*_*ik*_) is defined as

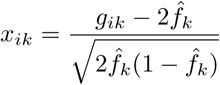

where 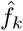 is the coded-allele frequency across the two samples. Effectively, the object containing the phenotypes is a three dimensional array, of size 11, 696 ×500 ×5. Under this data-generating process, the unconditional phenotypic variance (i.e., when it is not known to which population a given observation belongs) is given by

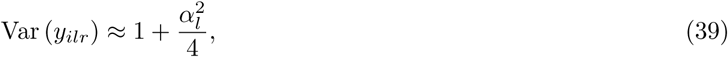

where the last term is the variance accounted for by stratification. Hence, the proportion of phenotypic variance explained by stratification, set at level *α*_*l*_, is given by 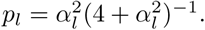

#### c.2 Three Populations

In the two-population design, we assigned one population a phenotypic mean of +*µ_l_* and the other a phenotypic mean of −*µ*_*l*_, where *µ*_*l*_ = *α*_*l*_/2. This amount of stratification explains proportion 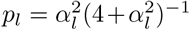 of the phenotypic variance. To keep the framework consistent, in the three-population design, in each run we assign one randomly selected population a mean of 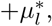 another randomly selected population a mean of 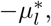 and the remaining population a mean of zero, where 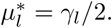 In this scenario, proportion

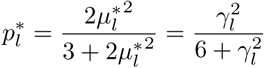

of the phenotypic variance is explained by 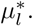 As we aim to have stratification explaining the same amount of phenotypic variance in both simulation designs, we require

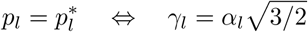

with *α*_*l*_ as set in the design for two populations. Consequently, we simulate data for three populations as follows:

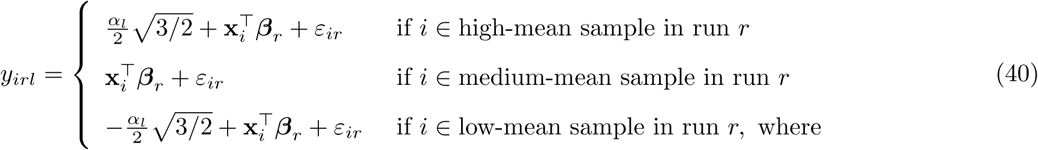

>

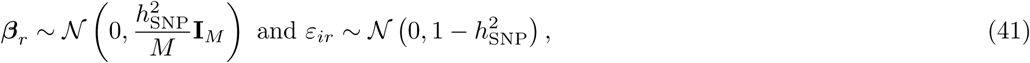

where all notation is the same as for two populations. In each run, the STR, HRS, and RS sample are randomly assigned to either have the low, medium, or high phenotypic mean. The object containing the phenotypes is a three dimensional array for the three-population simulation, of size 17, 544 ×500 ×5.

#### C.3 Empirical Analyses

In the empirical work, in the case additional stratification is added, the phenotype (e.g., height) for individual *i* (denoted by *y*_*i*_) is adjusted as follows:

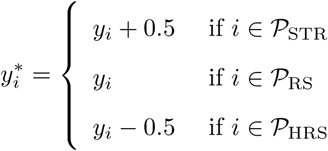

In the baseline analyses, *y*_*i*_ is used as phenotype, whereas in the design with additional stratification, 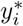 is used.

### D Fast GREML with Principal Components as Only Covariates

In case GREML estimation is used when only a subset of PCs is included as fixed-effects covariates, the computational complexity of REML can be reduced strongly. Following the origingal GREML model^9^, we have that

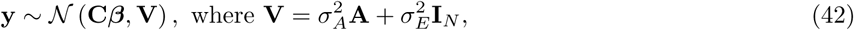

where **C** denotes the set of *K* leading principal components, from the *N*-by-*N* GRM **A** with eigendecomposition 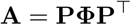 (i.e., **C** consists of the first *K* columns of **P**). Changing notation slightly, we consider the following expressions for the log-likelihood, gradient, and average information (AI) matrix, in line with standard GREML^9^:

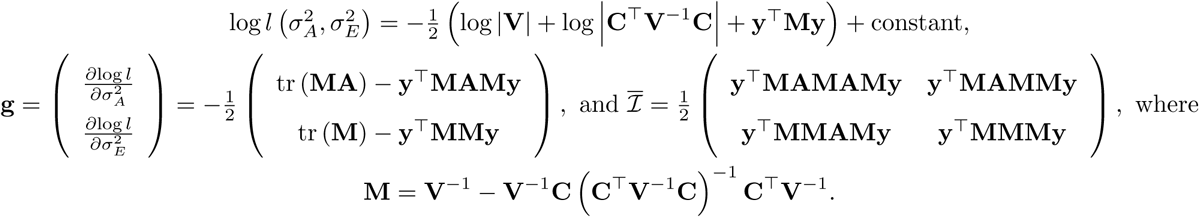

Using properties of the eigendecomposition, we have that

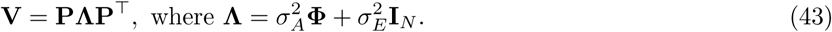

Importantly, **Λ** is a diagonal matrix and is therefore easily inverted. Further use of properties of eigendecompositions, and the fact that **C** is merely a subset of the columns of **P**, it follows that 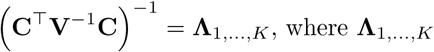 is a diagonal submatrix of **Λ**, containing the *K* largest values of **Λ**. Hence,

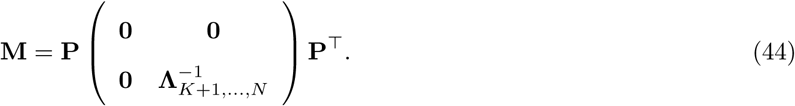

Defining 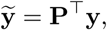 we can now rewrite the log-likelihood, gradient, and AI matrix as follows

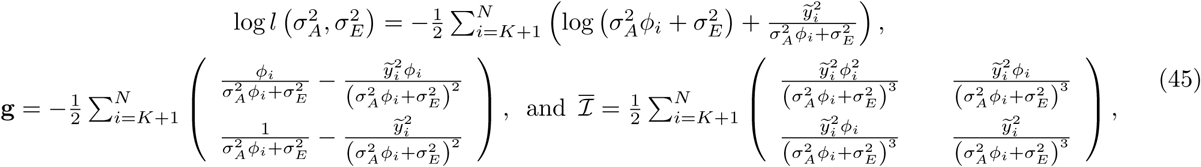

where *φ*_*i*_ denotes the *i*-th leading eigenvalue from the GRM and *y͂*_*i*_ denotes the *i*-th element of **y͂**. Noticing (i) that **y͂** consists only of *N* elements, which do not change over iterations, (ii) the same holds for the eigenvalues *φ*_*i*_, and (iii) the log-likelihood, gradient, and AI matrix are computationally easy functions of the variance components, the eigenvalues, and *y͂*_*i*_, it readily follows that AI-REML estimation (e.g., optimized using Newton's method) is computationally easy. In this specific model, the generalized least squares estimator is given by

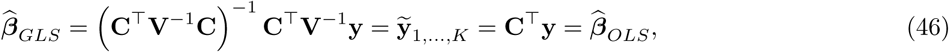

where 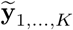 denotes the column vector containing the first *K* elements of 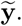.

In our simulation study with three subsamples we perform 2,500 separate REML analyses (i.e., 5 levels of stratification with 500 runs per level), with *N* ≈ 18k in each analysis. Exploiting the fact that the same GRM is used in all analyses and using our efficient algorithm, we carry out these GREML analyses in less than three minutes on a machine with 64GB RAM and 24 cores with a clock rate of 2.4GHz. Importantly, far less than the available 64GB RAM is used in these analyses.

## Acknowledgements

**Dutch national e-infrastructure** This work was carried out on the Dutch national e-infrastructure with the support of SURF Cooperative.

**HRS (Health and Retirement Study)** The HRS is sponsored by the National Institute on Aging (grant number NIA U01AG009740) and is conducted by the University of Michigan. The genotyping was funded separately by the National Institute on Aging (RC2 AG036495, RC4 AG039029). The genotyping was conducted by the NIH Center for Inherited Disease Research (CIDR) at Johns Hopkins University. Genotyping quality control and final preparation of the data were performed by the Genetics Coordinating Center at the University of Washington. Genotype data can be accessed via the database of Genotypes and Phenotypes (dbGaP, http://www.ncbi.nlm.nih.gov/gap, accession number phs000428.v1.p1). Researchers who wish to link genetic data with other HRS measures that are not in dbGaP, such as educational attainment, must apply for access from HRS. See the HRS website (http://hrsonline.isr.umich.edu/gwas) for details.

**RAND HRS** RAND HRS Data, Version N. Produced by the RAND Center for the Study of Aging, with funding from the National Institute on Aging and the Social Security Administration. Santa Monica, CA (September 2014). Researchers who wish to use the RAND HRS data need to register via the RAND website (http://www.rand.org/labor/aging/dataprod/hrs-data.html).

**RS (Rotterdam Study)** The generation and management of GWAS genotype data for the RS is supported by the Netherlands Organisation of Scientific Research NWO Investments (nr. 175.010.2005.011, 911-03-012). This study is funded by the Research Institute for Diseases in the Elderly (014-93-015; RIDE2), the Netherlands Genomics Initiative (NGI)/Netherlands Organisation for Scientific Research (NWO) project nr. 050-060-810. We thank Pascal Arp, Mila Jhamai, Marijn Verkerk, Lizbeth Herrera and Marjolein Peters for their help in creating the GWAS database, and Karol Estrada and Maksim V. Struchalin for their support in creation and analysis of imputed data. The RS is funded by Erasmus Medical Center and Erasmus University, Rotterdam, Netherlands Organization for the Health Research and Development (ZonMw), the Research Institute for Diseases in the Elderly (RIDE), the Ministry of Education, Culture and Science, the Ministry for Health, Welfare and Sports, the European Commission (DG XII), and the Municipality of Rotterdam. The authors are grateful to the study participants, the staff from the RS and the participating general practitioners and pharmacists. Researchers who wish to use data of the RS must obtain approval from the Rotterdam Study Management Team. They are advised to contact the PI of the RS, Dr. Arfan Ikram.

**STR (Swedish Twin Registry)** The Jan Wallander and Tom Hedelius Foundation (P2015-0001:1), the Ragnar Söderberg Foundation (E9/11), The Swedish Research Council (421-2013-1061; M-2205-1112), GenomEUtwin (EU/QLRT-2001-01254; QLG2-CT-2002-01254), NIH DK U01-066134, The Swedish Foundation for Strategic Research (SSF). The STR is financially supported by Karolinska Institutet. We wish to thank the Biobank at Karolinska Institutet for professional biobank service. Researchers interested in using STR data must obtain approval from the Swedish Ethical Review Board and from the Steering Committee of the Swedish Twin Registry. Researchers using the data are required to follow the terms of an Assistance Agreement containing a number of clauses designed to ensure protection of privacy and compliance with relevant laws. For further information, contact Patrik Magnusson.

**Individual acknowledgements** R.d.V. acknowledges funding from an ERC consolidator grant (647648 EdGe, 570 awarded to Philipp D. Koellinger). P.M.V. acknowledges support from the Australian National Health and Medical Research Council (grants 1078037 and 1113400). All the authors acknowledge valuable feedback provided by Philipp D. Koellinger, Naomi R. Wray, Michael E. Goddard, Matthew R. Robinson, Jian Yang, and Michel G. Nivard.

## Web Resources

`LDSC`^4^: github.com/bulik/ldsc

`GCTA`^9^: cnsgenomics.com/software/gcta/

`PLINK`^10,11^: www.cog-genomics.org/plink2

